# Cancer Phenotypic Plasticity Quantification using Morphology-Migration Coupled Metric in Live Label-Free Optical Microscopy

**DOI:** 10.64898/2026.07.06.736717

**Authors:** Saylee Muley, Krishna Agarwal, Biswajoy Ghosh

**Affiliations:** Department of Physics and Technology, UiT - The Arctic University of Norway, Tromsø, Norway

## Abstract

Cancer phenotypic plasticity drives invasion, treatment resistance, and relapse. Quantifying how cells dynamically couple morphology and migration in real time, without molecular labels, remains unsolved. Static molecular markers report on protein expression state rather than functional migratory behavior. Existing image-based metrics treat shape and migration as independent features, missing the coordinated coupling that defines plastic migratory states. We introduce Directional Shape Coupling (DSC), a quantitative metric purpose-built for live label-free imaging. DSC integrates movement direction consistency, shape deformation, and directional-shape alignment into a single interpretable score. Component weights are derived from PCA, adapting automatically to any dataset without manual tuning. Applied to differential interference contrast imaging of pancreatic cancer cells on a tissue-mimicking substrate recapitulating desmoplastic tumor stroma, DSC exhibited a large phenotype-associated effect size, *ε*^2^ = 0.65, across five distinct migratory phenotypes within a genetically homogeneous population, demonstrating that behavioral heterogeneity is structured and non-genetic. DSC encodes information orthogonal to classical shape and motion descriptors. Critically, DSC reveals that dynamic shape adaptation to mechanical cues rather than directional commitment drives phenotypic identity in this system. DSC provides the label-free imaging community a transparent, generalizable framework for quantifying dynamic non-genetic plasticity directly from live imaging data.

## Introduction

Phenotypic plasticity in cancer, the capacity of tumor cells to dynamically switch between distinct behavioral states without genetic change, is a key driver of invasion, treatment resistance, and relapse^1–3^. In pancreatic ductal adenocarcinoma (PDAC), this plasticity is shaped by a dense desmoplastic stroma that is among the stiffest of all malignancies, actively driving mechanoadaptive cell behavior through constitutively active oncogenic signaling^4–7^. Cells do not migrate persistently along fixed trajectories but continuously remodel their shape in response to local mechanical cues, adopting transient migratory states that are directly relevant to treatment escape^2,8,9^. Capturing this behavior quantitatively, in real time, without molecular labels, and with biological interpretability, remains a fundamental unmet need.

Current approaches fail to meet this need in complementary ways. Molecular profiling of epithelial-to-mesenchymal transition (EMT) markers, including epithelial markers such as E-cadherin and occludin, mesenchymal markers such as N-cadherin and fibronectin, and cytoskeletal regulators, provides snapshots of transcriptional or protein state but not of functional migratory behavior^10–12^. A cell population may express a mesenchymal signature while remaining collectively static, or conversely exhibit coordinated migratory plasticity with minimal molecular reprogramming^13–16^. Live-cell imaging offers direct observation of adaptive behavior^17–23^, but existing image-based metrics treat cell shape and migration as independent readouts, missing the coordinated coupling between them that defines plastic migratory states^24–27^. Integration of shape and motion has been explored using machine learning and deep learning^28,29^, but these approaches rely on complex non-linear models that function as black boxes, limiting interpretability and obscuring the biological mechanisms that drive plasticity^30,31^. Label-free imaging modalities such as differential interference contrast microscopy are particularly underserved, providing rich morphodynamic information across long timescales without phototoxicity or perturbation, yet no interpretable metric exists to extract coupled shape-motion behavior from these data^32,33^.

Here we introduce Directional Shape Coupling (DSC), a quantitative metric purpose-built for live label-free imaging^34,35^ that integrates real-time morphological remodeling with directional migration into a single interpretable score. DSC combines directional persistence, shape deformation, and directional-shape alignment using data-driven Principal Component Analysis (PCA)-derived weights, ensuring the metric reflects the dominant axes of phenotypic variance in any given dataset without manual parameter tuning. Applied to differential interference contrast imaging of MIA PaCa-2 PDAC cancer cells on a tissue-mimicking substrate recapitulating the stiffness of desmoplastic PDAC stroma, DSC accounts for a large phenotype-associated effect size (*ε*^2^ = 0.65) across manually annotated morphodynamic phenotypes within a genetically homogeneous population, encodes information orthogonal to classical shape and motion descriptors, and reveals that shape adaptation rather than directional persistence is the dominant mode of plasticity in this mechanically constrained system. DSC offers the label-free imaging community a transparent, adaptable framework for quantifying dynamic non-genetic plasticity directly from live imaging data, with broad implications for studying adaptive resistance across cancer types and microenvironmental conditions.

## Results

### Cell shape and migration dynamics uncover phenotypic states along the EMT continuum

Live-cell Differential Interference Contrast (DIC) imaging of MiaPaca-2 cells seeded in an inhouse substrate revealed pro-nounced heterogeneity in cell morphology and migratory behavior within the same experimental conditions (Figure 1, a-f). Representative fields illustrate the coexistence of multiple distinct morphological states, while accompanying trajectory analyses highlight corresponding diversity in migration patterns across cells (Figure 1, g-k). Based on qualitative inspection of cell shape, size, cell–cell adhesion, and apparent motility, cells were manually grouped into five distinct morphological types (Types A–E). The emergence of these distinct morphological states was consistently observed as a substrate-driven phenomenon across multiple independent experiments using this substrate^**?**^. Table 1 describes morphological and migratory characteristics of each of the cell types in detail. This qualitative classification highlights the extent of phenotypic diversity present during migration and provided the basis for subsequent quantitative analysis aimed at objectively capturing dynamic differences in migratory plasticity.

**Table 1.** Qualitative morphodynamic features used to define phenotypic cell types *A–E*. Morphological and migratory characteristics of phenotypic cell types *(A–E)* were assessed by manual observation of representative live-cell imaging data. Reported numeric values are approximate visual estimates and should be interpreted as descriptive ranges supporting phenotypic classification, not as automated or statistically quantified measurements. P.L., protrusion length; P.C., protrusion count.

| Type | Shape | Size ($\mu\text{m}$) | Cell–cell adhesion | Motility ($\mu\text{m}/\text{frame}$) | P. C. | P. L. ($\mu\text{m}$) |
| --- | --- | --- | --- | --- | --- | --- |
| A | Spherical, clustered | $25 \pm 5$ | High | $4 \pm 3$ | $\leq 1$ | $\leq 2$ |
| B | Elongated, spindle-like | $40 \pm 10$ | Low | $30 \pm 10$ | $2 \pm 2$ | $40 \pm 10$ |
| C | Small, rounded | $15 \pm 5$ | Minimal | $3 \pm 1$ | $\leq 3$ | $\leq 2$ |
| D | Irregular, spread-out | $50 \pm 10$ | Low | $20 \pm 10$ | $7 \pm 3$ | $50 \pm 20$ |
| E | Elongated, fusiform | $50 \pm 10$ | Low | $10 \pm 3$ | $3 \pm 2$ | $40 \pm 20$ |

**Figure 1.**
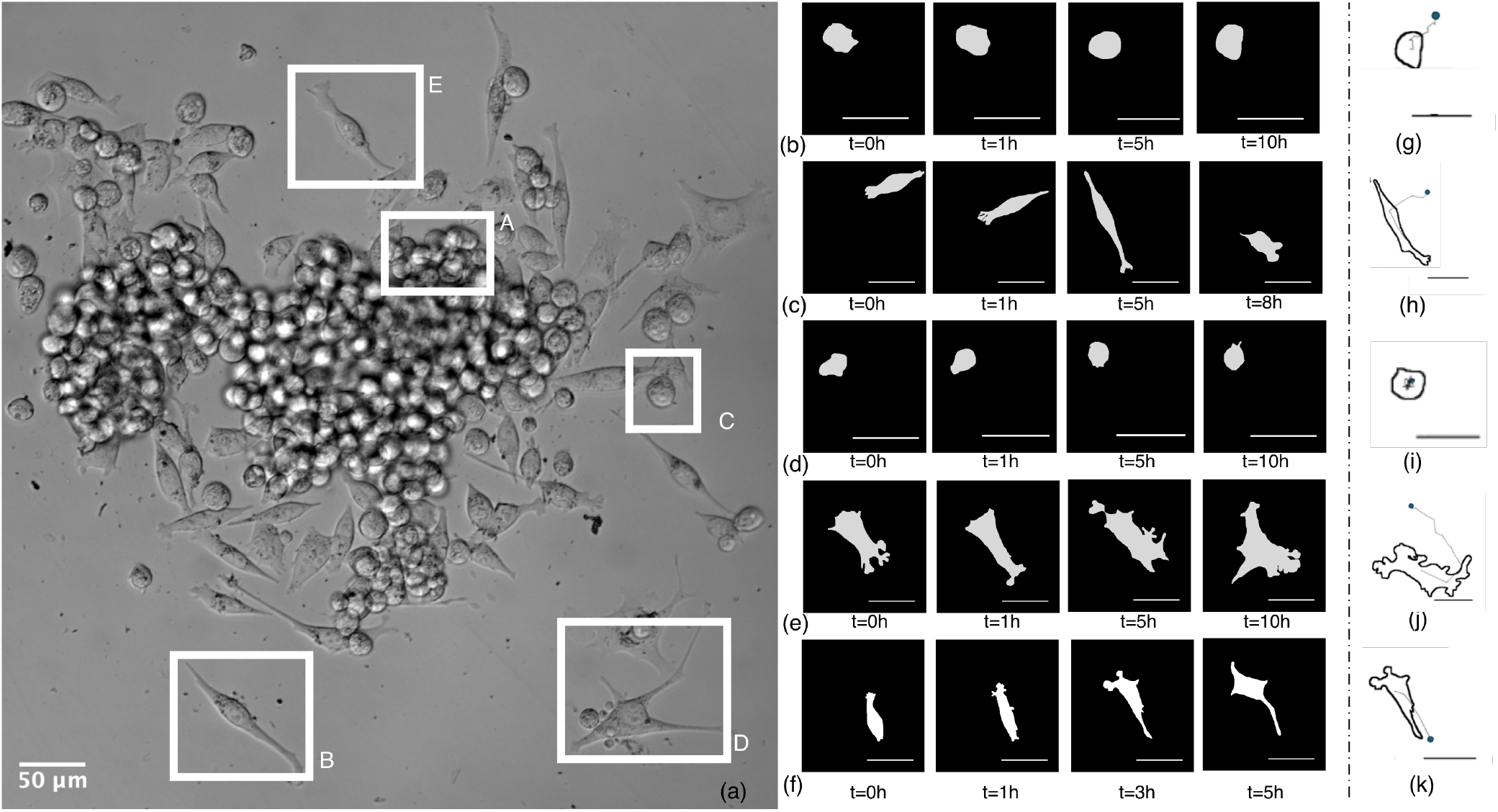
Morphodynamic heterogeneity reveals distinct phenotypic states in migrating MIA PaCa-2 PDAC cells. Live-cell imaging revealed pronounced morphodynamic heterogeneity among migrating MIA PaCa-2 cells, with cells displaying distinct combinations of morphology, shape evolution, and migration trajectory. These features were used to qualitatively classify cells into five phenotypic categories, *(A–E)*, representing different states across an EMT-associated plasticity spectrum. **a**, Differential interference contrast image acquired 8 h after seeding, showing heterogeneous cell morphologies within a single field of view. Representative cells assigned to categories *(A–E)* are highlighted. Scale bar, 50*µ*m. **b-f**, Temporal evolution of cell morphology for representative cells using binary outlines from each phenotypic category: type *A* in **(b)**, type *B* in **(c)**, type *C* in **(d)**, type *D* in **(e)**, type *E* in **(f)**. Scale bars, 50*µ*m. **g-k**, Representative migration trajectories corresponding to the same phenotypic categories shown in **b–f**, respectively. Each trajectory originates from the blue dot and follows the indicated path, illustrating differences in migration directionality, displacement and path geometry between phenotypic categories. Black scale bars, 50*µ*m. These characteristics are further explained in Table 1.

### Directional Shape Coupling formalizes morphodynamic heterogeneity into a quantitative cell-state metric

Quantifying the qualitative behaviors observed above enables a robust and objective characterization of cell migratory plasticity. While existing metrics quantify directional persistence or shape deformation independently, they often fail to capture their coupled temporal evolution or lack biomechanistic interpretability^28,29,36–38^. Motivated by the coexistence of distinct morphological states and migration patterns (Figure 1), we introduce a composite metric termed Directional Shape Coupling (DSC), designed to quantify how changes in cell shape and migration direction evolve together over time.

We define the DSC at time *t* as

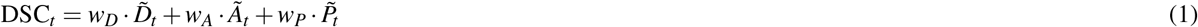

The DSC_*t*_ is defined as a weighted linear combination of three dimensionless features that capture distinct aspects of cell migratory behavior, namely deformation, directional alignment, and persistence. The deformation 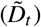 quantifies the extent to which a cell deviates from its resting shape. The directional alignment (*Ã*_*t*_) represents the degree to which cell motion is aligned between shape change and migration direction, while the persistence 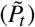 characterizes the consistency of cell movement over time, reflecting how steadily a cell maintains its migration direction. Each component is standardized using z-scoring, such that 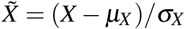 , ensuring comparability across features. These normalized variables are combined using dimensionless weighting coefficients (*w*_*D*_, *w*_*A*_, *w*_*P*_), which control the relative contribution of each feature to the final score. The resulting composite metric DSC_*t*_ is expressed in arbitrary units (a.u.), as it represents a weighted combination of normalized features rather than a directly measurable physical quantity.

Cell trajectories and shape features were extracted from TrackMate-generated XML files using a custom processing pipeline, yielding time-resolved descriptors of cell position, morphology, and protrusive activity.

We next describe each component of the DSC metric in detail, outlining how deformation, directional alignment and persistence together capture complementary features of cell morphodynamic plasticity.

#### Directional Persistence 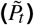

Directional persistence quantifies the temporal consistency of cell migration by measuring how long a cell maintains movement along a similar direction, while also accounting for the magnitude of displacement. For each cell, the centroid position at time *t* is denoted by *c*_*t*_, and the frame-to-frame displacement vector is defined as

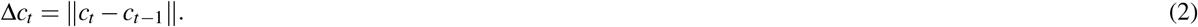

The instantaneous migration angle *θ*_*t*_ is computed from this displacement vector as

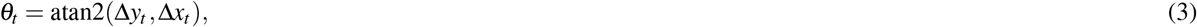

where Δ*x*_*t*_ and Δ*y*_*t*_ are the horizontal and vertical components of Δ*c*_*t*_, respectively. The continuous angle *θ*_*t*_ is then discretized into one of *N*_bins_ angular bins, yielding a direction index *d*_*t*_. Here, *N*_bins_ is a user-defined hyperparameter that determines the angular resolution used to classify migration direction; larger values provide finer directional resolution, whereas smaller values group a broader range of angles into the same direction class.

To quantify how long a cell maintains the same discretized migration direction, we defined the directional dwell length *l*_*t*_ as the number of consecutive frames up to time *t* for which the direction index remains unchanged:

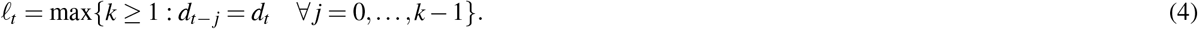

##### Algorithm 1

Directional Shape Coupling (DSC) computation

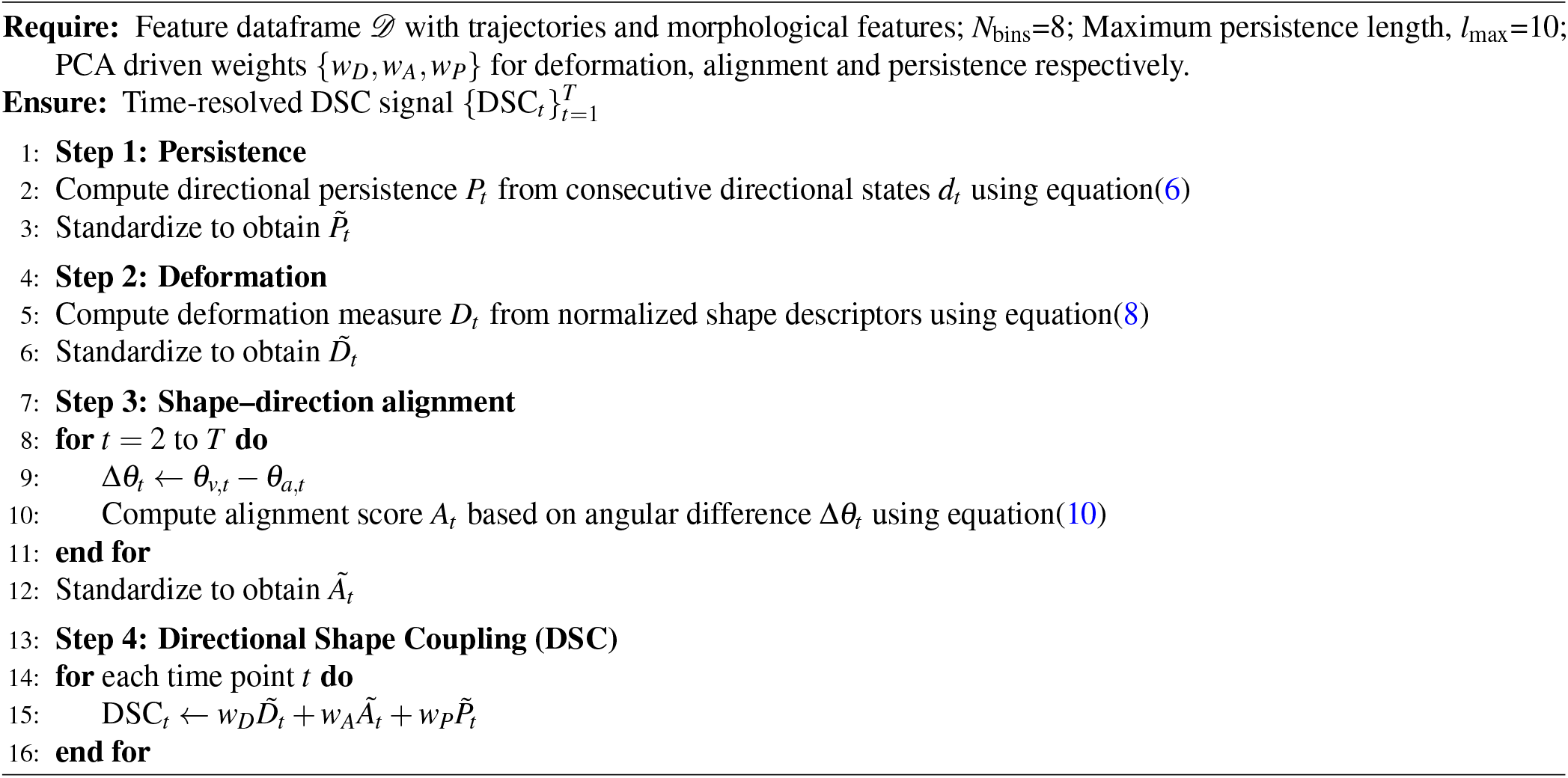

The dwell length is converted into directional continuity factor, *C*_*t*_, by normalizing it with respect to a maximum persistence threshold *l*_max_:

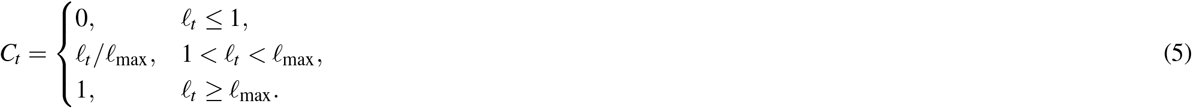

The parameter *l*_max_ is a predefined hyperparameter that sets the number of consecutive frames required for persistence saturation. Thus, short directional dwell events receive low scores, whereas trajectories that maintain the same direction for at least *l*_max_ consecutive frames are assigned the maximum persistence score. This saturation prevents very long uninterrupted directional segments from disproportionately dominating the metric.

To incorporate the extent of cell movement, the directional persistence score is weighted by the instantaneous step size, yielding the effective persistent displacement:

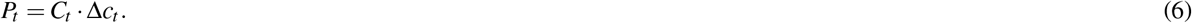

Here, *P*_*t*_ captures both directional consistency and displacement magnitude: cells that move far while maintaining a stable direction receive higher values than cells that move weakly or frequently change direction.

Finally, the persistence component used in the Directional Shape Coupling metric is obtained by z-score standardization:

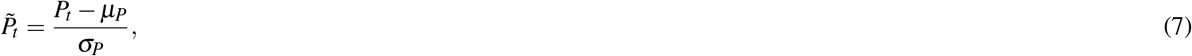

where *µ*_*P*_ and *σ*_*P*_ denote the mean and standard deviation of *P*_*t*_ across the analysed cell trajectories and time points. The resulting quantity 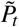 is dimensionless and represents the standardized persistence contribution to the DSC metric.

#### Shape Deformation (*D*_*t*_)

Shape deformation quantifies the extent to which a cell deviates from its resting morphological state and acquires elongated, protrusive or branched structures over time. At each time point *t*, deformation is computed from a set of global shape, skeletonization-derived and branching features extracted from the binary cell mask. Each feature is first standardized by z-score normalization, and the final deformation index is defined as the equally weighted average of all standardized features.

For a given feature *k*, the raw feature value *F*_*tk*_ was transformed into a z-score as

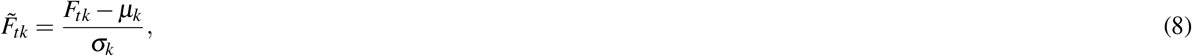

where *µ*_*k*_ and *σ*_*k*_ denote the mean and standard deviation of feature *k* across the analysed cells and time points. The deformation index is then calculated as

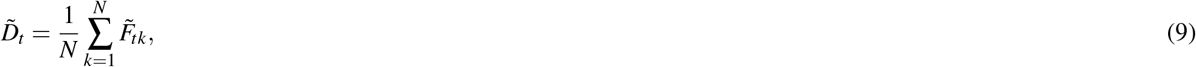

where *N* is the total number of deformation-associated features. Thus, each feature contributes equally to *D*_*t*_ after standardization, ensuring that no single descriptor dominates because of its original scale or unit.

The feature set included standard global shape descriptors and skeletonization-derived structural descriptors. Global shape features comprised shape index, area, perimeter, solidity, circularity, radius and aspect ratio, capturing overall cell size, compactness, elongation and boundary complexity. Skeletonization-derived features were included to capture protrusive architecture and internal shape topology that are not fully represented by global shape descriptors alone. Table 2 groups all deformation-associated features by feature class, with each class representing a distinct structural aspect of cell morphology. The table further defines how global shape and skeletonization-derived descriptors contribute complementary information to the deformation index and support phenotypic discrimination.

**Table 2.** Global shape and skeletonization-derived features used to compute the deformation index. The deformation index *D*_*t*_ was calculated from standardized global shape descriptors and skeletonization-derived structural descriptors. Global descriptors quantify overall cell size, compactness, elongation and boundary complexity. Skeletonization-derived descriptors quantify medial-axis structure, protrusive tips, branching density, branch extent and branch-size heterogeneity. These feature classes capture both large-scale cell morphology and fine protrusive architecture. Each feature was z-score standardized and contributed equally to the deformation index.

| Feature class | Feature | Definition | Biological interpretation / role in $D_t$ |
| --- | --- | --- | --- |
| **A. Global shape descriptors** |  |  |  |
| Cell size | Area | Projected area of the segmented cell mask | Cell spreading and overall cell size |
| Cell boundary | Perimeter | Boundary length of the segmented cell mask | Boundary extent and contour irregularity |
| Shape compactness | Circularity | Similarity of the cell outline to a circle | Compactness versus irregular or protrusive morphology |
| Shape compactness | Solidity | Ratio of mask area to convex hull area | Boundary indentation and non-convex protrusive structure |
| Shape complexity | Shape index | Perimeter normalized by cell area | Boundary complexity relative to cell size |
| Cell scale | Radius | Effective radius derived from the cell mask | Approximate cell size independent of boundary complexity |
| Elongation | Aspect ratio | Ratio of major to minor axis length | Cell elongation and morphological polarization |
| **B. Skeletonization-derived structural descriptors** |  |  |  |
| Skeleton extent | Skeleton pixels | Number of pixels in the skeletonized cell mask | Total medial-axis extent of the cell |
| Skeleton extent | Normalized skeleton length | Skeleton length normalized by cell size | Size-adjusted elongation and protrusive extension |
| Protrusive tips | Endpoints | Number of terminal nodes in the skeleton | Number of protrusive tips or terminal extensions |
| Skeleton topology | Connected components | Number of disconnected skeleton components | Continuity or fragmentation of the medial-axis structure |
| Branching complexity | Number of branches | Number of skeleton branch segments | Degree of protrusive branching |
| Branching complexity | Branch lengths | Lengths of individual skeleton branch segments | Distribution of protrusive extension lengths |
| Branching complexity | Branch density by area | Number of branches normalized by cell area | Branching complexity relative to cell size |
| Branching complexity | Branch density by length | Number of branches normalized by skeleton length | Branching frequency along the medial axis |
| Branch size | Mean branch length | Average length of branch segments | Typical protrusion or branch extension length |
| Branch size | Maximum branch length | Longest branch segment length | Dominant protrusive extension |
| Branch size | Total branch length | Sum of all branch segment lengths | Overall extent of branched protrusive architecture |
| Branch heterogeneity | Branch length standard deviation | Standard deviation of branch segment lengths | Heterogeneity in protrusion or branch sizes |

The skeletonization and branching features were particularly important for distinguishing phenotypic cell types with similar global morphology but different protrusive organization. Representative examples from each phenotypic class show substantial variation in skeletal architecture, including differences in skeleton length, terminal endpoints, branching density and branch-length heterogeneity (Figure 2). These features therefore provided critical discriminatory information for identifying highly protrusive, elongated or branched morphologies.

**Figure 2.**
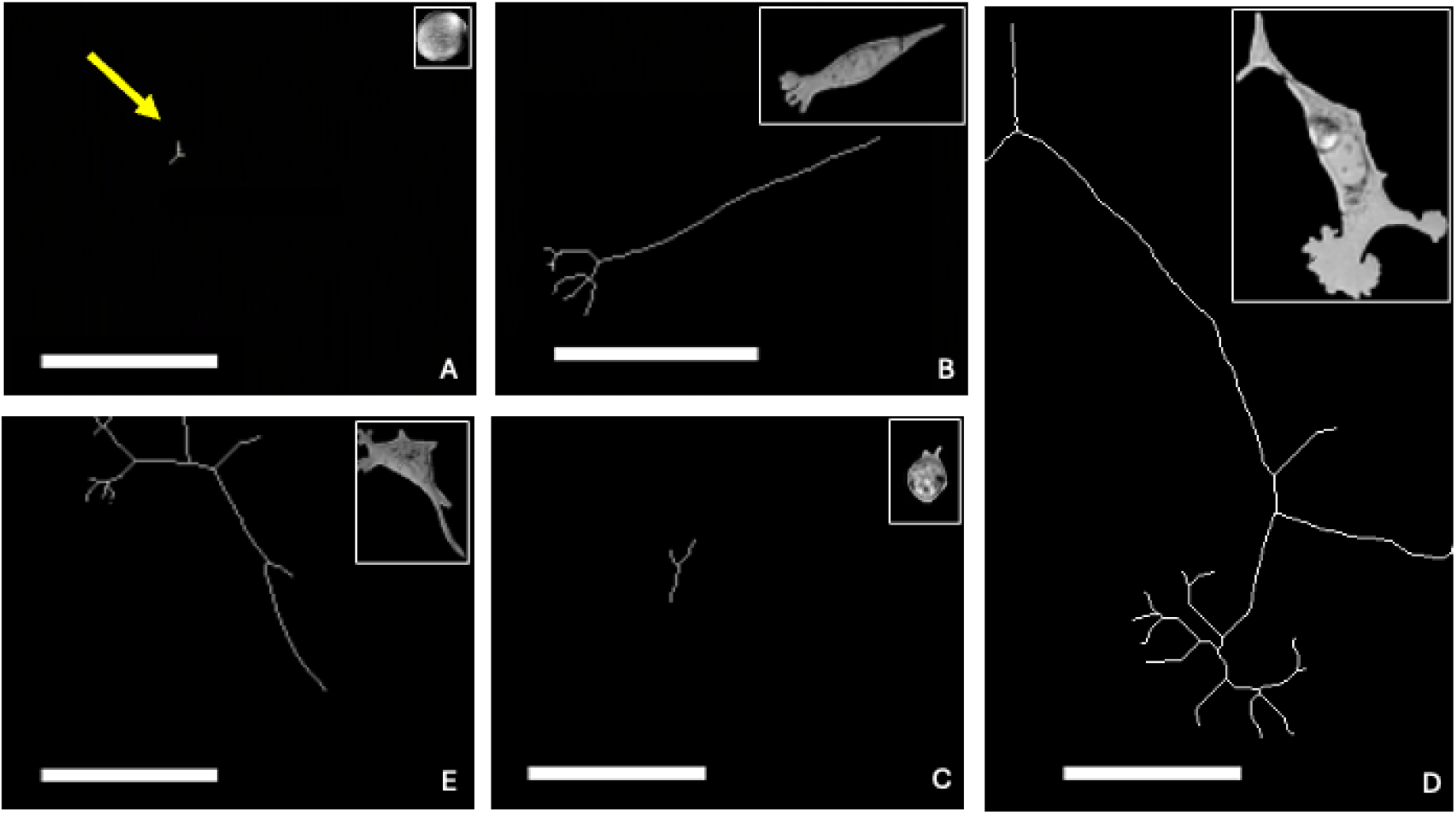
Skeletonization exposes structural heterogeneity underlying deformation-based phenotypic classification. Representative examples of phenotypic cell types *A–E* are shown as binary cell masks with their corresponding skeletonized representations. Skeletons were extracted as one-pixel-wide medial-axis structures from segmented cell masks, preserving key topological features of cell morphology. Across phenotypic classes, skeletons reveal marked differences in protrusive extent, elongation, endpoint number and branching complexity. These visual differences provide direct evidence that skeletonization-derived descriptors capture structural features not fully represented by global shape measurements alone, supporting their use as major contributors to the deformation index *D*_*t*_. Insets show the original segmented cells. Scale bars, 50 *µ*m.

Biologically, the deformation index represents a composite measure of morphological remodeling associated with changes in cytoskeletal organization, membrane protrusion and contractility. However, deformation alone does not indicate whether shape changes are aligned with productive migration. Therefore, deformation is integrated with directional alignment and persistence within the DSC framework to quantify coupled shape–migration plasticity.

#### Directional-Shape Alignment (*A*_*t*_)

Directional-shape alignment captures the instantaneous correspondence between the direction of migration and the principal orientation of cell shape. It is defined as

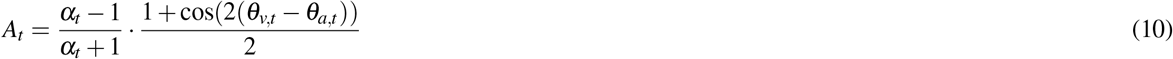

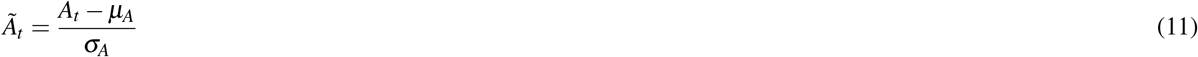

where, *α*_*t*_ = aspect ratio of ellipse, *θ*_*v,t*_ = cell velocity direction, *θ*_*a,t*_ = major axis direction

At each time point, segmented cell outlines are analyzed using TrackMate^39^ to obtain an ellipse approximation, from which the major axis orientation, *θ*_*a,t*_ and aspect ratio are extracted. The instantaneous migration direction, *θ*_*v,t*_ is computed from centroid displacement. Alignment is defined using a cosine similarity between the migration direction and the major axis orientation, scaled by the cell aspect ratio. The use of cos(2 (*θ*_*v,t*_ −*θ*_*a,t*_)), accounts for the axial symmetry of the ellipse, ensuring that orientations differing by 180^°^ are treated equivalently. Aspect-ratio weighting suppresses spurious alignment in near-isotropic cells, for which a principal axis is poorly defined. High values of *A*_*t*_ occur when elongated cells migrate along their major axis, indicating coordinated shape polarization and motion, whereas low values arise when deformation is orthogonal to migration or when shape anisotropy is weak. Alignment therefore serves as the key coupling term linking morphology and directionality.

### PCA-derived weighting integrates deformation, alignment and persistence into DSC

After defining the deformation, directional alignment and persistence components, z-score normalization is performed to place all three quantities on a comparable dimensionless scale. Principal component analysis (PCA) was then applied to the standardized component space to determine the relative contribution of each component to the dominant axis of morphodynamic variability. Component weights were obtained from the normalized absolute loadings of the first principal component, allowing the metric to emphasize the dominant source of morphodynamic variation without manually tuning the relative component weights. The resulting PCA-derived weights were *w*_*D*_ = 0.39 for deformation, *w*_*A*_ = 0.40 for directional alignment and *w*_*P*_ = 0.21 for persistence. The final DSC score was calculated as a weighted linear combination of the standardized components as shown in eq. 1.

Thus, for each cell at each time point, DSC integrates deformation, directional alignment and persistence into a single dimensionless score. The PCA-derived weighting indicated that deformation and alignment contributed similarly to the dominant variation in the dataset, whereas persistence contributed a smaller fraction. This weighting procedure allowed the DSC metric to retain interpretability while adapting the relative contribution of each morphodynamic component to the measured dataset.

High DSC values therefore correspond to time points at which standardized deformation, alignment and persistence jointly contribute positively to the composite score, whereas lower DSC values indicate weaker combined contribution of these components. The resulting DSC time series provides a unified quantitative representation of morphodynamic coupling and is subsequently used to compare phenotypic cell states.

### Directional Shape Coupling distinguishes migratory phenotypes

To assess whether DSC distinguishes migratory phenotypes, we compared the distributions of deformation, directional alignment, persistence and DSC across the five phenotypic cell types. Metrics were evaluated at the cell-time-point level, where each data point corresponds to one tracked cell observed at one imaging frame. In total, the analysis included *n*_cells_ = 135 tracked cells and *N*_timepoints_ = 14,326 cell-time points. The temporal spacing between consecutive observations was determined by the live-cell imaging frame rate. The number of cell-time points per phenotypic class was *N*_*A*_ = 5,146, *N*_*B*_ = 1,909, *N*_*C*_ = 2,960, *N*_*D*_ = 3,263 and *N*_*E*_ = 1,048, corresponding to 35.9%, 13.3%, 20.7%, 22.8% and 7.3% of the total dataset, respectively.

Because the number of observations differed across phenotypic classes, we used rank-based non-parametric statistical tests for group comparisons. The Kruskal–Wallis test and Mann–Whitney U test are appropriate for comparing groups without assuming normally distributed measurements, and Kruskal–Wallis-type analyses can be applied to groups with unequal sample sizes because the test is based on ranks rather than raw values^40–42^. The observed class imbalance was moderate, with an approximately 4.9-fold difference between the largest and smallest phenotypic classes, and therefore did not invalidate the use of these non-parametric tests. Group sizes are nevertheless reported explicitly to make the sampling structure transparent and to support interpretation of statistical power and effect estimates, particularly for the least represented phenotype.

Box-and-whisker plots showed that the individual DSC components exhibited partial overlap between phenotypic classes, whereas the integrated DSC score showed stronger separation across cell types, including differences in both median values and distributional spread (Figure 3a). This indicates that combining deformation, alignment and persistence into a single weighted metric improves discrimination of migratory phenotypes compared with any individual component alone.

**Figure 3.**
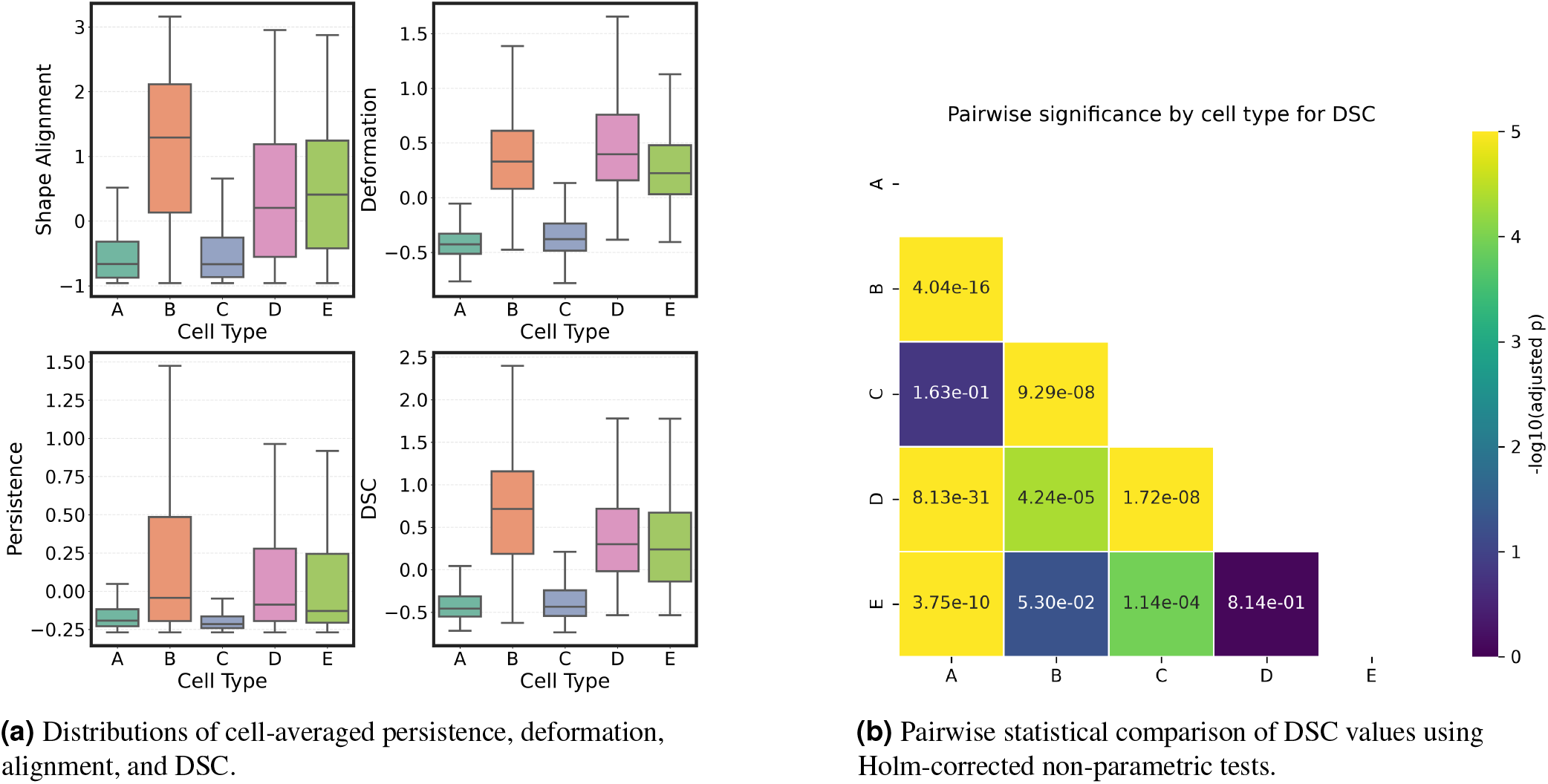
Directional Shape Coupling captures phenotype-dependent variation in morphodynamic behaviour. **a**, Distributions of frame-resolved shape alignment, deformation, persistence and Directional Shape Coupling (DSC) values across phenotypic cell types *A–E*. Each data point corresponds to one cell-time point, defined as one tracked cell at one imaging frame. Box plots show the median, interquartile range and overall spread of each metric for each phenotypic class. Individual components show different degrees of phenotype-dependent separation: deformation and shape alignment display strong shifts between several cell types, whereas persistence shows greater overlap and weaker separation. The integrated DSC score combines these component-level behaviours and provides a unified morphodynamic readout, with clear separation between several classes. For example, types B and E show more distinct median DSC values than in deformation alone, although their broad distributional overlap indicates partially shared morphodynamic behaviour. **b**, Pairwise statistical comparison of DSC distributions between phenotypic cell types using Mann–Whitney U tests with Holm correction for multiple comparisons. Values within the matrix indicate Holm-adjusted *p*-values for each pairwise comparison. The color scale represents −log_10_(*p*_adj_), where larger values and brighter colors indicate smaller adjusted *p*-values and therefore stronger statistical evidence for phenotype-dependent separation. Several phenotype pairs, including *A–B, A–D, A–E, B–C, C–D* and *C–E*, show strong separation based on DSC. In contrast, pairs such as *A–C* and *D–E* show weak or no significant separation, consistent with overlapping DSC distributions and shared morphodynamic characteristics. The *B–E* comparison is borderline after correction, indicating that although these phenotypes differ visually in DSC distribution, their overlap limits statistical separation at the corrected significance threshold.

To evaluate the statistical robustness of phenotype-dependent differences in DSC, pairwise comparisons were performed between all cell types using Mann–Whitney U tests with Holm correction for multiple comparisons. The resulting pairwise significance matrix showed that several phenotype pairs remained significantly different after correction, whereas others showed weaker or non-significant separation (Figure 3b). This pattern is consistent with the observed overlap in DSC distributions and suggests that some phenotypic classes share partially overlapping morphodynamic behaviours.

The omnibus Kruskal–Wallis test further confirmed significant differences in DSC across the five phenotypic groups.

DSC showed a strong phenotype-dependent effect, with *H* = 192, *p <* 10^−40^ and *ε*^2^ = 0.655. The effect size indicates that phenotypic group explained approximately 65.5% of the rank-based variability in DSC values. These results show that DSC provides a statistically robust and practically meaningful descriptor of morphodynamic phenotype at frame-level temporal resolution (Table 3). Although deformation produced the largest overall effect size, it failed to distinguish phenotypically similar elongated populations (e.g. Types *B* and *E* see Figure S3). Incorporating alignment and persistence resolved these differences, demonstrating that the additional discriminatory power arises from coupling rather than deformation alone.

**Table 3.** Integrated DSC provides stronger phenotype-dependent separation than individual morphodynamic components. Frame-resolved deformation, alignment, persistence and DSC values were compared across phenotypic cell types *A–E* using Kruskal–Wallis tests. Each observation corresponds to a track-level summary. The analysis included *n*_cells_ = 135 tracked cells and *N*_timepoints_ = 14,326 cell-time points. Effect size was reported as epsilon squared, *ε*^2^, representing the proportion of rank-based variability in each metric associated with phenotypic class. Deformation showed the largest phenotype-dependent effect, indicating that shape remodeling is a major discriminator of cell type, while DSC retained strong separation by integrating deformation with alignment and persistence into a unified morphodynamic metric.

| Metric | $H$ | $p$-value | $\epsilon^2$ |
| --- | --- | --- | --- |
| Deformation, $\tilde{D}_t$ | 214 | $< 10^{-45}$ | 0.73 |
| Alignment, $\tilde{A}_t$ | 146 | $< 10^{-30}$ | 0.49 |
| Persistence, $\tilde{P}_t$ | 35 | $< 10^{-7}$ | 0.11 |
| Directional Shape Coupling, DSC | 192 | $< 10^{-40}$ | 0.65 |

### Comparison between vimentin intensity and DSC across cell states

To relate the morphodynamic cell-state metric to an established cytoskeletal marker, we compared DSC with vimentin intensity across representative cells from phenotypic classes *A–E*. Vimentin was used because it is a type III intermediate filament protein and a commonly used marker of mesenchymal/EMT-associated cytoskeletal organization, with increased vimentin expression linked to cytoskeletal remodeling and migratory phenotypes^43,44^.

Immunofluorescence imaging showed heterogeneous vimentin intensity and organization across the MIA PaCa–2 cell population, with representative cells (*n* = ≈10) from phenotypic classes *A–E* displaying distinct combinations of vimentin signal and cell morphology (Figure 4). Quantification of mean vimentin intensity and mean DSC showed that the two measurements did not vary identically across phenotypic classes (Figure 5). Type *B* showed high vimentin intensity and high DSC, with mean vimentin intensity of approximately 149 a.u. and mean DSC of approximately 0.55. Type *D* also showed high DSC, approximately 0.47, despite a lower mean vimentin intensity of approximately 132 a.u. In contrast, type *C* showed the highest mean vimentin intensity, approximately 164 a.u., but low DSC, approximately −0.14.

**Figure 4.**
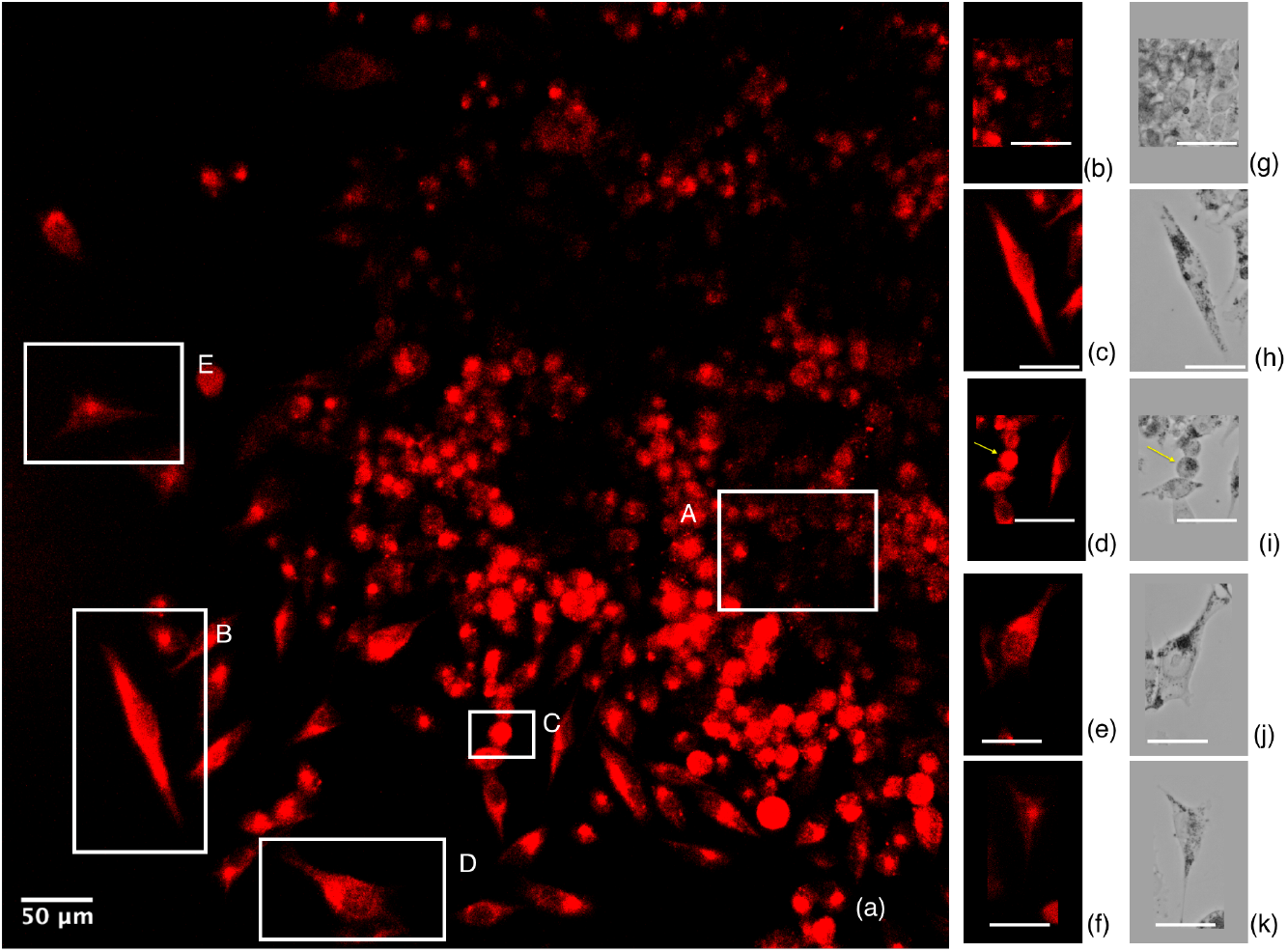
Vimentin immunofluorescence reveals cytoskeletal heterogeneity across phenotypic cell states. **a**, Representative vimentin immunofluorescence image of MIA PaCa-2 PDAC cells showing heterogeneous vimentin intensity and organization within the same field of view. Vimentin is used as a EMT-associated cytoskeletal marker and is linked to cytoskeletal remodeling during migratory cell-state transitions. Representative cells corresponding to phenotypic types *A–E* are indicated by white boxes. Scale bar, 50 *µ*m. **b–f**, Magnified vimentin fluorescence images of representative cells from phenotypic classes *A–E*, respectively, showing differences in vimentin signal intensity, spatial organization and cell-associated cytoskeletal morphology. **g–k**, Corresponding phase contrast images of the same representative cells shown in **b–f**, illustrating the associated cell morphology. The panel demonstrates that phenotypic classes differ not only in overall cell shape but also in the organization and abundance of vimentin-positive cytoskeletal structures. These representative examples motivate the quantitative comparison between vimentin intensity and Directional Shape Coupling shown in Figure 5. Scale bars, 50 *µ*m.

**Figure 5.**
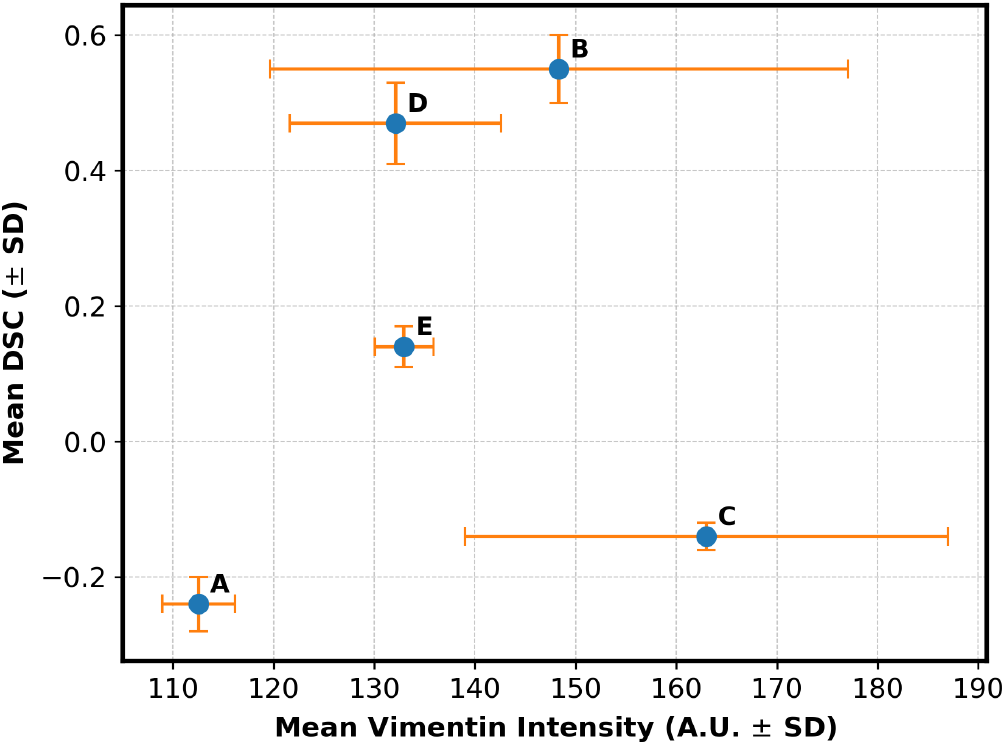
Mean vimentin intensity and Directional Shape Coupling vary differently across phenotypic cell types. Mean vimentin intensity was plotted against mean DSC for phenotypic cell types *A–E*. Points indicate class means and error bars indicate standard deviation. Type *B* showed high vimentin intensity and the highest DSC, whereas type *C* showed the highest vimentin intensity but low DSC. Types *D* and *E* had similar vimentin intensities but different DSC values, indicating that the two measurements vary differently across phenotypic states.

Differences between phenotypes were also evident among classes with similar vimentin intensities. Types *D* and *E* showed comparable mean vimentin intensities of approximately 132–133 a.u., but differed in mean DSC, with type *D* at approximately 0.47 and type *E* at approximately 0.14. Similarly, types *B* and *C* both showed elevated vimentin intensity relative to type *A*, but their DSC values differed strongly, with type *B* showing high DSC and type *C* showing low DSC. These measurements show that phenotypic classes with similar or elevated vimentin intensity can display different DSC values, indicating that vimentin intensity and DSC capture non-equivalent features of the classified cell states. The purpose of this comparison is not to validate DSC against vimentin, but to demonstrate that DSC captures complementary functional information that is not reflected by static marker intensity alone.

### Relationship between DSC and dynamic-only migration metrics

To determine whether DSC merely recapitulates existing dynamic descriptors of motion, we performed principal component analysis (PCA)^45^ combining DSC metrics (mean and variability) with commonly used dynamic-only measures, including velocity autocorrelation^26^, direction autocorrelation^46^, turning angle variance^47^, and MSD-derived persistence length^38^.

In the motion-only PCA (Table 4), the first three principal components explained 91.6% of the total variance, indicating that classical migration descriptors occupied a relatively low-dimensional feature space. PC1 explained 40.8% of the variance and was dominated by direction autocorrelation and turning angle variance, representing a straight-versus-reorienting migration axis. PC2 explained 28.7% of the variance and was dominated by velocity autocorrelation, whereas PC3 explained 22.1% of the variance and was primarily associated with MSD-derived persistence length. Thus, the motion-only feature space mainly captured directional memory, reorientation dynamics and trajectory-scale persistence.

**Table 4.** Comparison of PCA feature loadings for motion-only versus joint DSC and dynamic motion feature spaces. PC1 and PC2 loadings are shown for each feature. In the motion-only space, PC1 is dominated by directional motion metrics (direction_autocorr, turning_angle_variance), explaining 40.8% of variance. Upon inclusion of DSC features, the primary variance axis shifts to reflect shape plasticity (dsc_mean, dsc_std, alignment_mean), demonstrating that DSC introduces a novel and dominant source of biological variance that is not recoverable from motion metrics alone. Bold values indicate the strongest absolute contributors (≥ 0.40) to each principal component.

| Feature | Motion-Only PCA |  |  | Joint DSC + Motion PCA |  |  |
| --- | --- | --- | --- | --- | --- | --- |
|  | PC1 (40.8%) | PC2 (28.7%) | PC3 (22.1%) | PC1 (43.4%) | PC2 (17.8%) | PC3 (12.5%) |
| *DSC Features* |  |  |  |  |  |  |
| dsc_mean | — | — | — | **0.482** | 0.154 | -0.173 |
| dsc_std | — | — | — | **0.425** | 0.191 | 0.259 |
| alignment_mean | — | — | — | **0.439** | 0.076 | -0.262 |
| deformation_mean | — | — | — | 0.391 | 0.215 | -0.362 |
| persistence_mean | — | — | — | 0.298 | 0.120 | **0.596** |
| *Dynamic Motion Features* |  |  |  |  |  |  |
| velocity_autocorr | 0.036 | **0.814** | 0.478 | 0.008 | -0.386 | **-0.527** |
| direction_autocorr | **0.688** | 0.067 | 0.290 | -0.229 | **0.613** | -0.151 |
| turning_angle_variance | **-0.672** | 0.308 | 0.011 | 0.264 | **-0.578** | 0.202 |
| msd_persistence_length | -0.271 | -0.488 | **0.829** | 0.149 | 0.119 | 0.075 |
| **Variance Explained** | 91.6% (3 PCs) |  |  | 73.7% (3 PCs) |  |  |

When DSC-derived features were included, the structure of the PCA space changed. In the joint DSC + motion PCA, PC1 explained 43.4% of the variance and was dominated by DSC mean, DSC variability, alignment and deformation. In contrast, classical dynamic-only metrics contributed weakly to PC1, with velocity autocorrelation showing a near-zero loading. PC2 explained 17.8% of the variance and was mainly associated with direction autocorrelation and turning angle variance, indicating that reorientation-related motion features were retained on a separate axis. PC3 explained 12.5% of the variance and included stronger contributions from persistence mean and velocity autocorrelation.

These results show that DSC is not reducible to standard dynamic-only measures of migration. Instead, inclusion of DSC introduces a dominant shape–motion axis associated with coordinated deformation, alignment and morphodynamic variability, while classical motion metrics remain primarily associated with directional memory and reorientation dynamics.

## Discussion

Directional Shape Coupling (DSC) is introduced as a quantitative framework that integrates real-time morphological remodeling with directional migration, capturing a behavioral dimension of phenotypic plasticity^48^ that conventional analyses cannot resolve^10–12,24–26,28,29^. By validating DSC against manually identified migratory phenotypes in a genetically homogeneous pancreatic cancer cell population, we demonstrate that coupling between shape and direction provides discriminatory power beyond individual features, supporting the idea that coordinated migration states represent a distinct and clinically relevant axis of phenotypic adaptation (see Figure 3, Table 3). The dynamic-only migration metrics form a closed low-dimensional space describing directional memory and turning behavior, whereas inclusion of DSC reveals an orthogonal axis corresponding to coordinated morphodynamic organization (Table 4).

MIA PaCa-2 is a well-established model of pancreatic ductal adenocarcinoma, a malignancy characterized by exceptionally poor prognosis, dense desmoplastic stroma, and inherent resistance to standard chemotherapy including gemcitabine^4–7^. A defining feature of PDAC progression is the capacity of tumor cells to dynamically switch between epithelial and mesenchymal states, enabling invasion, dissemination, and post-treatment re-establishment^2,8,9^. Critically, this plasticity operates within genetically homogeneous populations, suggesting that behavioral heterogeneity is regulated at the level of phenotypic state rather than genetic mutation. Resistance to therapy in PDAC is increasingly understood as an adaptive rather than purely genetic phenomenon, driven by dynamic state transitions that enable subpopulations to survive therapeutic pressure and re-establish tumor growth^4–7^. The large phenotype-associated effect size observed for DSC, *ε*^2^ = 0.65, within such a genetically homogeneous population supports the conclusion that non-genetic behavioral heterogeneity is a structured and quantifiable phenomenon. (Table 3). This large phenotype-associated effect, achieved using a label-free imaging metric without molecular perturbation or genetic profiling, suggests that morphodynamic coupling encodes biologically meaningful information about cell state that is accessible directly from live imaging data.

The constitutive activation of MAPK, PI3K/AKT, and Wnt signaling in MIA PaCa-2 cells directly regulates cytoskeletal organization, membrane protrusion, cell polarity, and migratory behavior, processes that are reflected in the morphodynamic features quantified by DSC. This mechanistic connection between signaling pathway activity and morphodynamic output means that DSC is not merely measuring morphological noise but is instead reporting on the downstream functional consequences of oncogenic signaling at the single-cell level. The phenotypic spectrum observed across Types *A* through *E* therefore reflects the dynamic equilibrium of cells sampling different signaling states within a genetically homogeneous population, consistent with established models of non-genetic tumor heterogeneity driving adaptive resistance.

The comparison between DSC and vimentin expression (Figure 4 and 5) reveals an important dissociation. Type *C* cells display elevated vimentin intensity but low DSC, indicating that cytoskeletal priming for mesenchymal behavior does not necessarily translate into coordinated migratory execution. This finding suggests that vimentin abundance reflects molecular potential rather than functional plasticity, and that a dynamic metric capturing the real-time coupling of shape and motion provides information inaccessible to static molecular markers. DSC provides a quantitative measure consistent with functional execution of plasticity. This distinction has direct implications for how adaptive resistance is assessed, a cell may be molecularly primed for invasion while remaining functionally static, and DSC captures this difference where conventional markers cannot.

The data-driven weighting of DSC components (see page 7) offers a biological insight beyond metric construction. The relatively low weight of persistence compared to alignment and deformation reflects the hybrid migratory character of MIA PaCa-2 cells on a tissue-mimicking substrate. Rather than sustaining directional trajectories, these cells continuously remodel their shape in response to local mechanical cues, a mechanoadaptive behavior consistent with the heterogeneous stromal architecture of pancreatic ductal adenocarcinoma. This pattern aligns with current understanding that PDAC cells navigate rather than march through their microenvironment, with shape adaptation serving as the primary mode of environmental sensing. The emergence of five distinct morphological states as a substrate-driven phenomenon, consistently observed across independent experiments, further indicates that the mechanical microenvironment actively shapes the phenotypic landscape of the population.

The graded separation observed across phenotypic classes is itself a biological result. The proximity of types *B* and *E*, and the partial overlap of *A* and *C*, reflects a genuine continuum rather than discrete epithelial or mesenchymal endpoints (Figure 3, 5). This is consistent with current models of hybrid EMT states, in which cells occupy intermediate positions along a plasticity spectrum rather than switching between fixed programs. DSC captures gradations along this spectrum, providing a quantitative axis for positioning cells relative to one another without imposing artificial categorical boundaries. Importantly, the partial non-separation of certain class pairs does not represent a limitation of the metric but rather evidence that the EMT spectrum in this system is continuous and that DSC is sensitive enough to reflect that biological reality.

Among the three DSC components, deformation accounted for the largest phenotype-associated effect size overall, reflecting its sensitivity to the broad morphological diversity present across the five cell types (Table 3). However, this population-level dominance did not translate uniformly into pairwise discriminative power. Types *B* and *E*, both elongated and highly deformed, could not be significantly distinguished by deformation alone, with pairwise comparison yielding a non-significant result (see figure S3). This reveals a fundamental limitation of single-feature descriptors in populations where phenotypic diversity is not uniformly distributed – high overall variance does not guarantee resolution of morphologically similar but behaviorally distinct states. DSC increased separation between *B* versus *E* relative to deformation alone, by incorporating how deformation was coordinated with migration direction and directional consistency. This demonstrates that the discriminative power of DSC emerges specifically from the coupling between components rather than from any single dominant feature, and that the relevant behavioral differences between certain phenotypes are encoded in the relationship between shape and motion rather than in shape magnitude alone.

DSC reveals that migratory plasticity is intermittent and state-like rather than stable or monotonic, with individual cells transitioning between coordinated and decoupled shape-motion regimes over time. Cells exhibiting transient high-DSC episodes, briefly coordinating shape remodeling with directional motion before reverting, represent a subpopulation of particular clinical relevance (Supplementary Figure S2). These cells are not persistently invasive but are opportunistically plastic, adopting migratory behavior transiently in ways that are invisible to endpoint assays or static molecular profiling. This temporal dimension of plasticity may be especially consequential in the context of treatment escape and tumor re-establishment following therapy, where small subpopulations transiently accessing invasive states can seed relapse without constitutive commitment to a mesenchymal program.

The use of PCA-derived weights provides an unsupervised, data-adaptive weighting scheme rather than manually tuned component weights. In new datasets, the weights can be recalibrated to reflect the dominant axes of morphodynamic variation, supporting the use of DSC as an adaptable analysis framework rather than a fixed system-specific score. This adaptability directly addresses the single cell line scope of the current study and provides a principled path toward application across cancer types, drug treatment conditions, and microenvironmental contexts. Future work extending DSC to patient-derived organoids and co-culture systems will be critical for assessing whether morphodynamic coupling serves as a scalable readout of adaptive resistance potential linked to clinical outcomes.

This study is limited to a single PDAC cell line cultured on one mechanically defined substrate. Future studies should evaluate DSC across additional cell lines, primary patient-derived models, and drug perturbation experiments to establish broader applicability.

## Methods

The complete computational pipeline for DSC derivation is shown in Figure 6. Live-cell DIC image sequences were first preprocessed, followed by cell segmentation and trajectory tracking to obtain time-resolved cell masks and centroid positions. Global shape descriptors, motility features and skeletonization-derived structural features were then extracted from each cell at each frame. These features were used to calculate deformation, directional alignment and persistence, which were standardized, weighted using PCA-derived component weights and combined to generate the final Directional Shape Coupling score.

**Figure 6.**
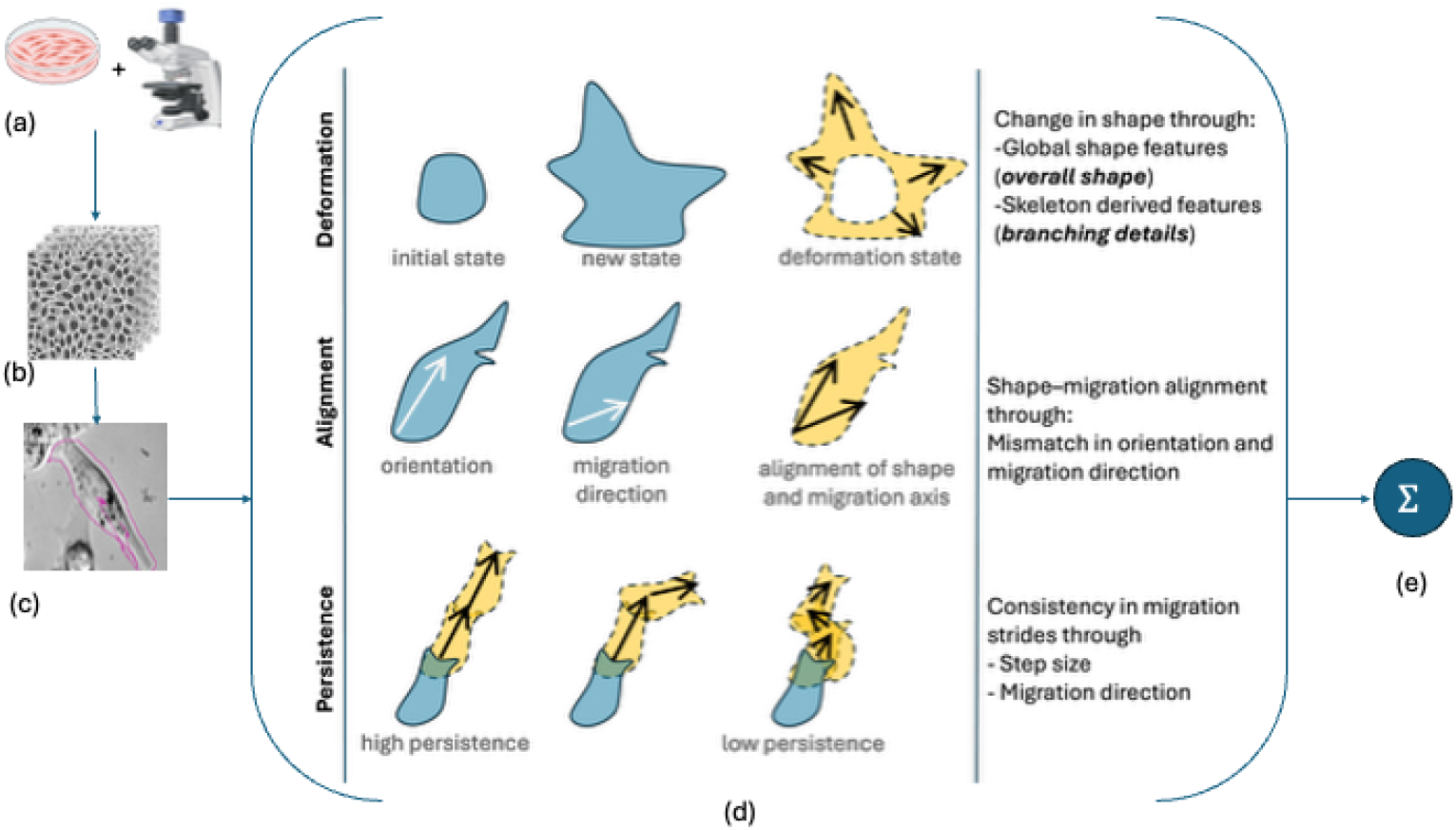
Conceptual and computational workflow for deriving Directional Shape Coupling from live-cell imaging. **a**, MIA PaCa-2 PDAC cells were cultured and imaged using live-cell microscopy to capture time-resolved changes in cell morphology and migration. **b**, Image processing to enhance cell segmentation and tracking **c**, Individual cells segmented and tracked to obtain binary cell masks, cell outlines and centroid trajectories for each frame. **d**, Directional Shape Coupling (DSC) formulated from three complementary morphodynamic components. Deformation quantifies changes in cell shape relative to the resting morphological state using global shape descriptors and skeletonization-derived features that capture protrusive and branching architecture. Alignment quantifies the correspondence between the cell shape axis and the migration direction, measuring whether morphological polarization is coordinated with movement. Persistence quantifies the temporal consistency of migration by combining step size with maintenance of migration direction across consecutive frames. **e**, The three standardized components were integrated using weighted summation to compute the final DSC score, providing a frame-resolved measure of coordinated shape remodeling and directional migration.

### Cell Culture

Experiments were performed using the MIA PaCa-2 pancreatic cancer cell line. Cells were cultured in Dulbecco’s Modified Eagle Medium (DMEM) supplemented with appropriate serum and growth factors and maintained at 37°C in a humidified incubator with 5% *CO*_2_. Cells were routinely passaged to ensure stable growth prior to seeding on hydrogels, following standard sterile cell culture procedures.

### Hydrogel Fabrication

Hydrogels were fabricated in-house using 10% (w/v) gelatin methacryloyl (GelMA) and photocrosslinking with lithium phenyl-2,4,6-trimethylbenzoylphosphinate (LAP) as photoinitiator at 365 nm UV exposure for 50 seconds, following a previously described protocol^49^. Multimodal mechanical characterization^50,51^ including compression testing, shear rheology, frequency-dependent storage modulus, and differential scanning calorimetry confirmed that 10% GelMA yields a stiffness in the range of 8-30 kPa, consistent with the desmoplastic stromal microenvironment of pancreatic ductal adenocarcinoma^49^. PDAC stroma is among the stiffest of all malignancies, with solid stress values approaching 10 kPa and tissue stiffness several magnitudes higher than healthy pancreatic tissue, driven by dense ECM deposition and stellate cell activation^52^. The mechanical properties of our substrate thus recapitulate the stiffened microenvironment under which MIA PaCa-2 cells exhibit adaptive migratory behavior in vivo. Following fabrication, hydrogels were equilibrated in culture medium and cells were seeded onto the hydrogel surface and allowed to adhere and spread before imaging.

### Selection of Cell-line

MIA PaCa-2 cells (Cytion, catalog number 300438) were used between passages 5 and 10 to ensure phenotypic stability. Cells were maintained in DMEM-high glucose supplemented with 10% FBS at 37°C in a humidified atmosphere with 5% CO2. This cell line carries homozygous KRAS p.Gly12Cys and CDKN2A deletion and constitutively activates MAPK, PI3K/AKT, and Wnt signaling pathways, producing intrinsic phenotypic heterogeneity under identical culture conditions without additional genetic perturbation

### Imaging Setup

Live-cell imaging was conducted using differential interference contrast (DIC) microscopy under controlled physiological conditions. Time-lapse imaging was performed for 20 hours, with images acquired every 5 minutes. Temperature was maintained at 37°C using a stage-top incubator to ensure cell viability throughout the experiment. All recordings were acquired at 20× magnification, providing sufficient resolution to capture dynamic changes in cell morphology and migration.

### Data Acquisition

Time-lapse sequences were acquired using an automated DIC imaging system optimized to minimize phototoxicity and focal drift during long-term imaging. Prior to each experiment, the system was calibrated to ensure consistent illumination, contrast, and focus stability. This setup generated image sequences suitable for quantitative analysis of both morphological remodeling and migratory behavior.

### Image Analysis and Cell Tracking

Image analysis was performed using FIJI^53^ in combination with custom Python^54^ scripts. Raw image sequences were first imported into FIJI, where standard preprocessing steps, including contrast enhancement and intensity normalization, were applied to ensure consistency across time-lapse recordings. Prior to DSC calculation, cells were manually classified into five phenotypic groups based on predefined morphological and migratory criteria summarized in Table 1. This qualitative classification served only as the reference grouping for evaluating DSC and was not used during feature extraction or metric computation. Cell segmentation was then carried out using Cellpose (advanced detection mode)^55,56^, enabling robust extraction of cell boundaries across a wide range of morphologies. The resulting segmented masks were used to obtain accurate contour representations for each cell at every time point. Cell tracking was performed in FIJI using the TrackMate framework^39^, which provides flexible integration of tracking algorithms. Trajectories were reconstructed using either a Linear Assignment Problem (LAP)–based tracker^57^, suitable for smooth and continuous motion, or a Kalman filter–based approach^58^ to handle intermittent or complex migration patterns. These complementary strategies enabled reliable reconstruction of cell trajectories across the full duration of the imaging sequence, minimizing track discontinuities during direction changes or transient shape fluctuations. Following segmentation and tracking, per-frame morphological features, trajectory data, and associated metadata were exported to Python for downstream analysis. Frame-wise DSC components were computed from these data, and group-level statistical inference was performed using track-level summary values, with each track treated as one independent observational unit. Unless otherwise stated, hyperparameter values of *N*_*bins*_ = 8 and *l*_*max*_ = 10 were selected to provide sufficient directional resolution while avoiding excessive sensitivity to frame-to-frame directional fluctuations.

## Acknowledgements

This work was supported by the MABIT-funded project (2024-0101 MarGel), UiT Talent funding, the H2020 FET Open RIA project OrganVision (Grant No.964800), and the H2020 ERC Starting Grant project 3D-nanoMorph (Grant No.804233). All results presented in this study were generated as part of the Cymoplive project.

## Data Availability

The data that support the findings of this study are available from the corresponding author upon reasonable request.

## Code Availability

The code that support the findings of this study are available from the corresponding author upon reasonable request.

## Competing interests

B.G. is the lead developer of Cymoplive, a pre-commercial cancer phenotypic profiling platform under development at UiT – The Arctic University of Norway, and is an inventor on patent application(s) related to technologies described in this work. The remaining authors declare no competing interests.

## Notes

### Competing Interest Statement

Biswajoy Ghosh is the lead developer of Cymoplive, a pre-commercial cancer phenotypic profiling platform under development at UiT The Arctic University of Norway, and is an inventor on patent application(s) related to technologies described in this work. The remaining authors declare no competing interests.

