## Supplementary Information for "Cancer Phenotypic Plasticity Quantification using Morphology-Migration Coupled Metric in Live Label-Free Optical Microscopy"

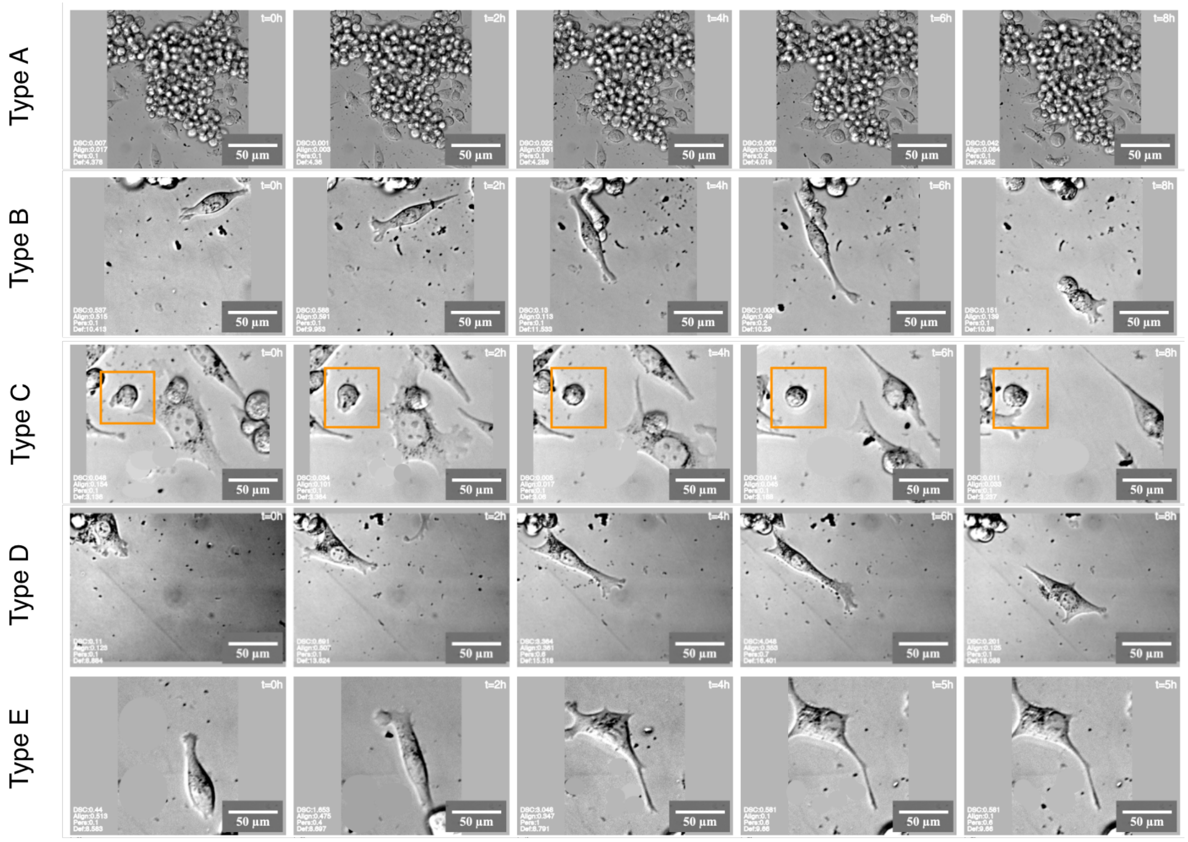


**Figure S1.** Time-resolved DIC morphodynamics with DSC overlays. Representative differential interference contrast (DIC) image sequences from five cell types from the same sample are shown as rows (top to bottom), with columns corresponding to elapsed time (left to right: t=0, 2, 4, 6, and 8h). Within each tile, the upper-right label indicates time, and the lower-left overlay reports frame-wise quantitative readouts: DSC, alignment, persistence, and deformation (dimensionless values, rounded to up to three decimal places). Together, the panel illustrates heterogeneous temporal behavior with the same sample, spanning dense multicellular organization and more isolated single-cell states, with corresponding variation in the overlaid DSC metrics.

**Table S1. Comparison of DSC component weights and phenotype-separation performance across weighting strategies.** For each Directional Shape Coupling (DSC) formulation, the weights assigned to deformation, directional alignment and persistence are shown together with the corresponding phenotype-separation statistics across cell types A–E. Overall group separation was assessed using the Kruskal–Wallis test, and effect size was reported as epsilon squared, $\varepsilon^2$, representing the proportion of rank-based variability in DSC associated with phenotypic class. Pairwise phenotype separation was assessed using Mann–Whitney U tests with Holm correction for multiple comparisons.

| **DSC formulation** | $w_D$ | $w_A$ | $w_P$ | $H$ | $p$**-value** | $\varepsilon^2$ |
|---|---|---|---|---|---|---|
| PCA-derived DSC | 0.39 | 0.40 | 0.21 | 192 | $< 10^{-40}$ | 0.655 |
| Equal-weight DSC | 0.33 | 0.33 | 0.33 | 191 | $< 10^{-40}$ | 0.654 |
| Deformation-emphasized DSC | 0.5 | 0.3 | 0.2 | 201 | $< 10^{-42}$ | 0.689 |
| Alignment-emphasized DSC | 0.3 | 0.5 | 0.2 | 183 | $< 10^{-38}$ | 0.623 |
| Persistence-emphasized DSC | 0.2 | 0.3 | 0.5 | 178 | $< 10^{-37}$ | 0.606 |

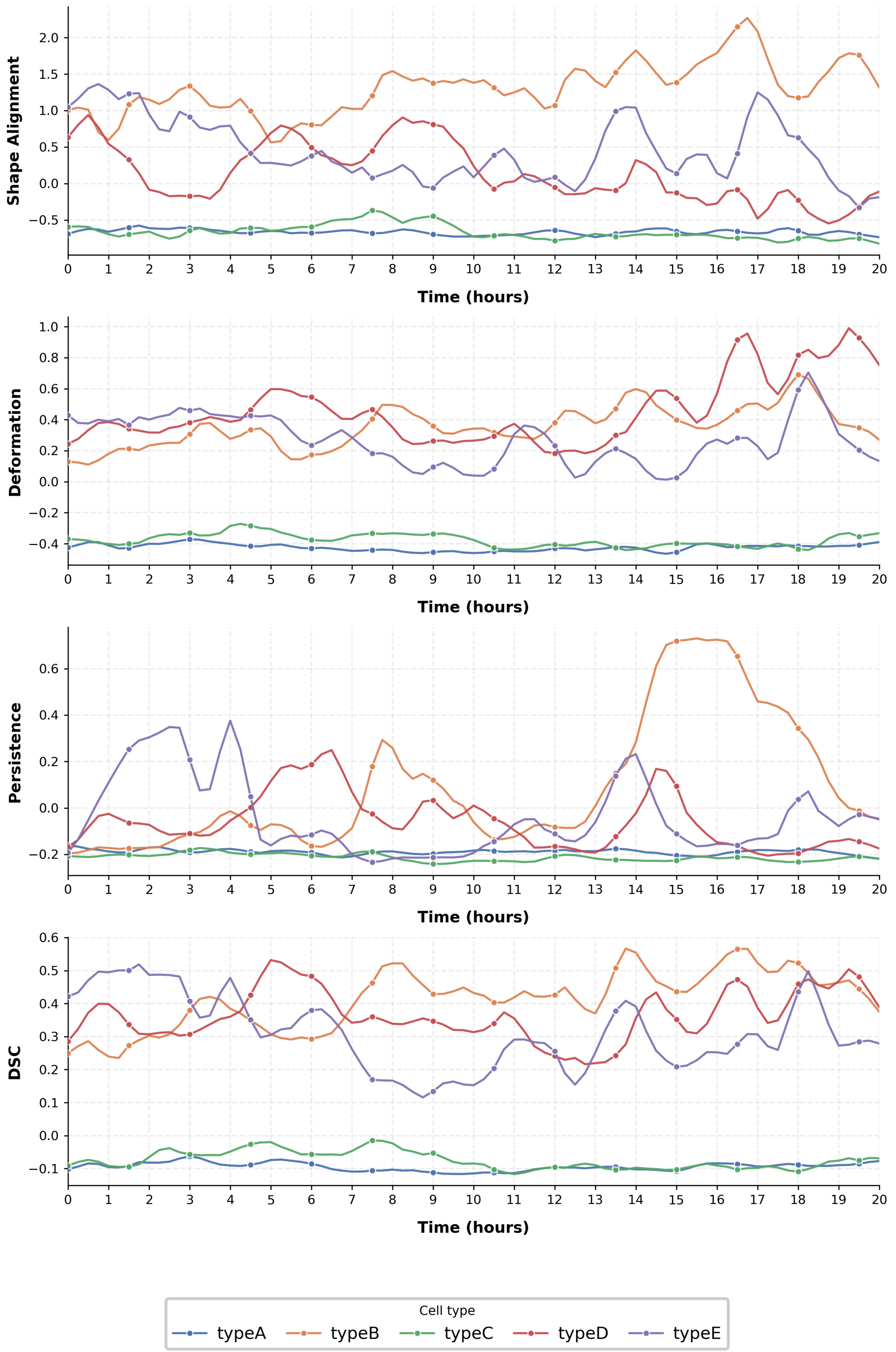


**Figure S2.** Temporal evolution of cell morphology and motility behavioral metrics. (a) Temporal evolution of cell morphological and migratory metrics across phenotypes (types A–E) over a 20-hour period. Line plots show dynamic changes in (top to bottom) shape alignment, deformation, persistence, and directional shape coupling (DSC). Distinct phenotypic behaviors are observed, with type B and D exhibiting higher variability and elevated values in alignment, deformation, and DSC, suggesting enhanced migratory coordination. In contrast, type A and C maintain consistently low values across all metrics, indicating limited motility and morphological changes.

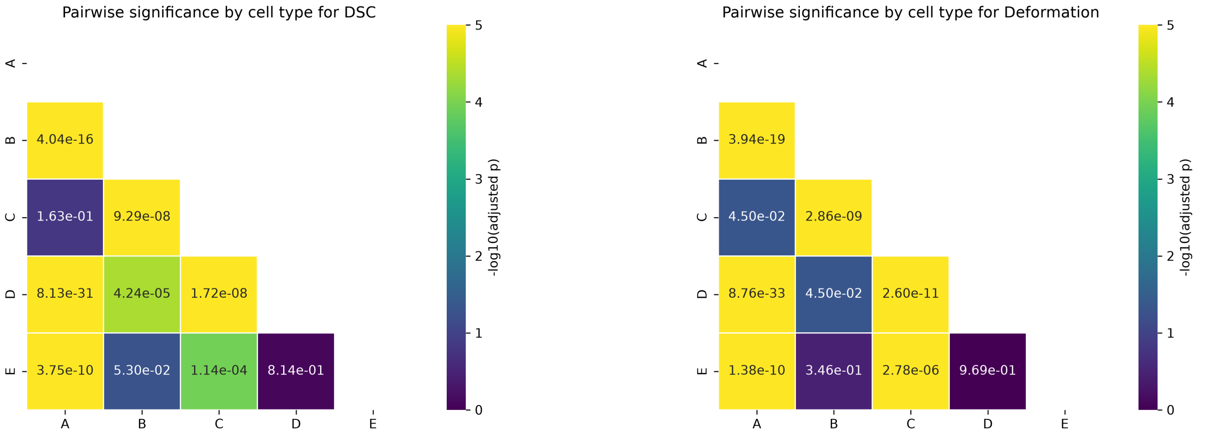


(a) (b)

**Figure S3. DSC and deformation show distinct pairwise separation patterns across phenotypic cell types.** Pairwise comparisons between phenotypic cell types A–E were performed using Mann–Whitney U tests with Holm correction. Heatmaps show adjusted $p$-values for **a,** Directional Shape Coupling (DSC) and **b,** deformation. Cell values indicate $p_{\text{adj}}$, and colors represent $-\log_{10}(p_{\text{adj}})$, where brighter colors indicate stronger statistical evidence for separation. The color scale was capped at 5. DSC showed strong separation for several phenotype pairs, including *A–B*, *A–D*, *A–E*, *B–C*, *C–D* and *C–E*. Deformation also showed strong separation for multiple pairs but failed to distinguish *B–E* and *D–E*. Notably, the *B–E* comparison was non-significant for deformation ($p_{\text{adj}} = 3.46 \times 10^{-1}$) and borderline for DSC ($p_{\text{adj}} = 5.30 \times 10^{-2}$), indicating that DSC increased pairwise separation relative to deformation alone. These results show that deformation captures broad phenotypic variance, whereas DSC provides a complementary integrated measure of shape–motion coupling that improves resolution for selected morphodynamically related phenotypes.
